# Selective germline genome edited pigs and their long immune tolerance in Non Human Primates

**DOI:** 10.1101/2020.01.20.912105

**Authors:** Lijin Zou, Youlai Zhang, Ying He, Hui Yu, Jun Chen, Delong Liu, Sixiong Lin, Manman Gao, Gang Zhong, Weicheng Lei, Guangqian Zhou, Xuenong Zou, Kai Li, Yin Yu, Gaofeng Zha, Linxian Li, Yuanlin Zeng, Jianfei Wang, Gang Wang

**Author notes:** Correspondence should be addressed to L.Z., Y.Y., L.L., Y.Z., J.W., G.W. These authors contributed equally to this work (Lijin Zou, Youlai Zhang). These authors contributed equally to this work (Guangqian Zhou, Xuenong Zou, Kai Li, Yin Yu, Linxian Li, Yuanlin Zeng, Jianfei Wang, Gang Wang).

## Abstract

Organ transplantation is the only curative treatment for patients with terminal organ failure, however, there is a worldwide organ shortage. Genetically modified pig organs and tissues have become an attractive and practical alternative solution for the severe organ shortage, which has been made possible by significant progress in xenotransplantation in recent years. The past several decades witnessed an expanding list of genetically engineered pigs due to technology advancements, however, the necessary combination of genetic modifications in pig for human organ xenotransplantation has not been determined. In the current study, we created a selective germline genome edited pig (SGGEP). The first triple xenoantigens (GGTA, B4GAL, and CAMH) knockout somatic cells were generated to serve as a prototype cells and then human proteins were expressed in the xenoantigen knockout cells, which include human complement system negative regulatory proteins (CD46, CD55, and CD59); human coagulation system negative regulatory proteins thrombomodulin (THBD); tissue factor pathway inhibitor (TFPI); CD39; macrophage negative regulatory proteins (human CD47); and natural killer cell negative regulatory human HLA-E. After the successful establishment of SGGEP by the nuclear tranfer, we engrafted SGGEP skin to NHP, up to 25 days graft survival without immunosuppressive drugs was observed. Because a pig skin graft does not impact the success of a subsequent allograft or autograft or vice versa, thus our SGGEP could have a great potential for clinical value to save severe and large area burn patients and the other human organ failure. Therefore, this combination of specific gene modifications is a major milestone and provides proof of concept to initiate investigator-initiated clinical trials (IITs) in severe burn patients with defined processes and governance measures in place and the other clinical application.

## Introduction

Organ transplantation is a major medical milestone that has saved the lives of 139,024 patients worldwide in 2017 with end stage organ failure (http://www.transplant-observatory.org/contador1/). However, the global donor shortage has limited its practical clinical application and further development. Genetically modified pigs have emerged as a potential alternative organ source, driving further research and development in xenotransplantation from pig to human(1-3).

Genetic modification of the pig is necessary to account for the differences between the pig and human genome, especially from the immune and molecular compatibility aspects. The utilization of CRISPR/Cas9 technology has resulted in the accelerated generation of more genetically modified pig lines (4). Nonetheless, determining which combination of genetic modification is necessary to launch a human clinical trial remains an open question (5). Currently, although some different multi-genes modified pigs are proposed or created, in vivo transplantation studies especially NHP experiments are yet to be carried to prove the effectiveness and safety of those genes to be modified or modified pigs. Meanwhile, the attention should be also paid to which patient populations in which disease areas could be early benefited from xenotransplantation (6).

Therefore, we adopted a strategy for developing a selective germline genome editied pig (SGGEP)(4, 7). First, triple xenoantigen (Glycoprotein alpha-galactosyltransferase 1, GGTA; β-1,4-N-acetyl-galactosaminyl transferase 2, B4GAL and, cytidine monophosphate-N-acetylneuraminic acid hydroxylase,CAMH) knockout somatic pig cells were generated to serve as a baseline for further gene editing. Then, with only one transfection step, we expressed a cluster human proteins including human complement system negative regulatory proteins (CD46, CD55, and CD59), human coagulation system negative regulatory proteins thrombomodulin (THBD), tissue factor pathway inhibitor (TFPI); CD39; macrophage negative regulatory proteins (human CD47); and natural killer cell negative regulatory human Human leukocyte antigen class I histocompatibility antigen, alpha chain E(HLA-E), thus, by using genome modification and cloning technologies, we created a preclinical and potentially clinical useful SGGEP.

After the successful production of SGGEP and the harvest of SGGEP-derived xenografts, we then performed xenotransplantation in several animal models, including NHP to test whether these genetic modifications combinations are justified for translational medicine applications and launching the clinical trial. Our results showed that SGGEP skin graft could survive functionally on NHP up to 25 days without the administration of immunosuppressive drugs. Considering that a pig skin graft does not affect the success of a subsequent allograft, or vice versa (8), therefore, this is a major milestone for skin xenotransplantation and serves as a proof of concept to initiate investigator-initiated clinical trials (IITs) in severe, life-threatening burn patients.

## Results

### Flow cytometric analysis of wildtype and SGGEP skin fibroblast phenotype confirmed SGGEP’s expected protein expression pattern

Flow cytometric analysis of SGGEP pig fibroblast and WT fibroblast confirmed the deletion of the three xenoantigens and the insertion of eight human genes. SGGEP fibroblasts and WT pig fibroblasts were collected for analysis. The fluorescence intensity of the three xenoantigens in the WT fibroblasts was notably increased compared to that of the isotype control (Fig.1a) indicating high xenoantigens expression. Nearly complete deletion of the three xenoantigens in the SGGEP fibroblasts is shown by no shift in fluorescence intensity compared to an isotype control (Fig. 1b).

**Figure 1.**
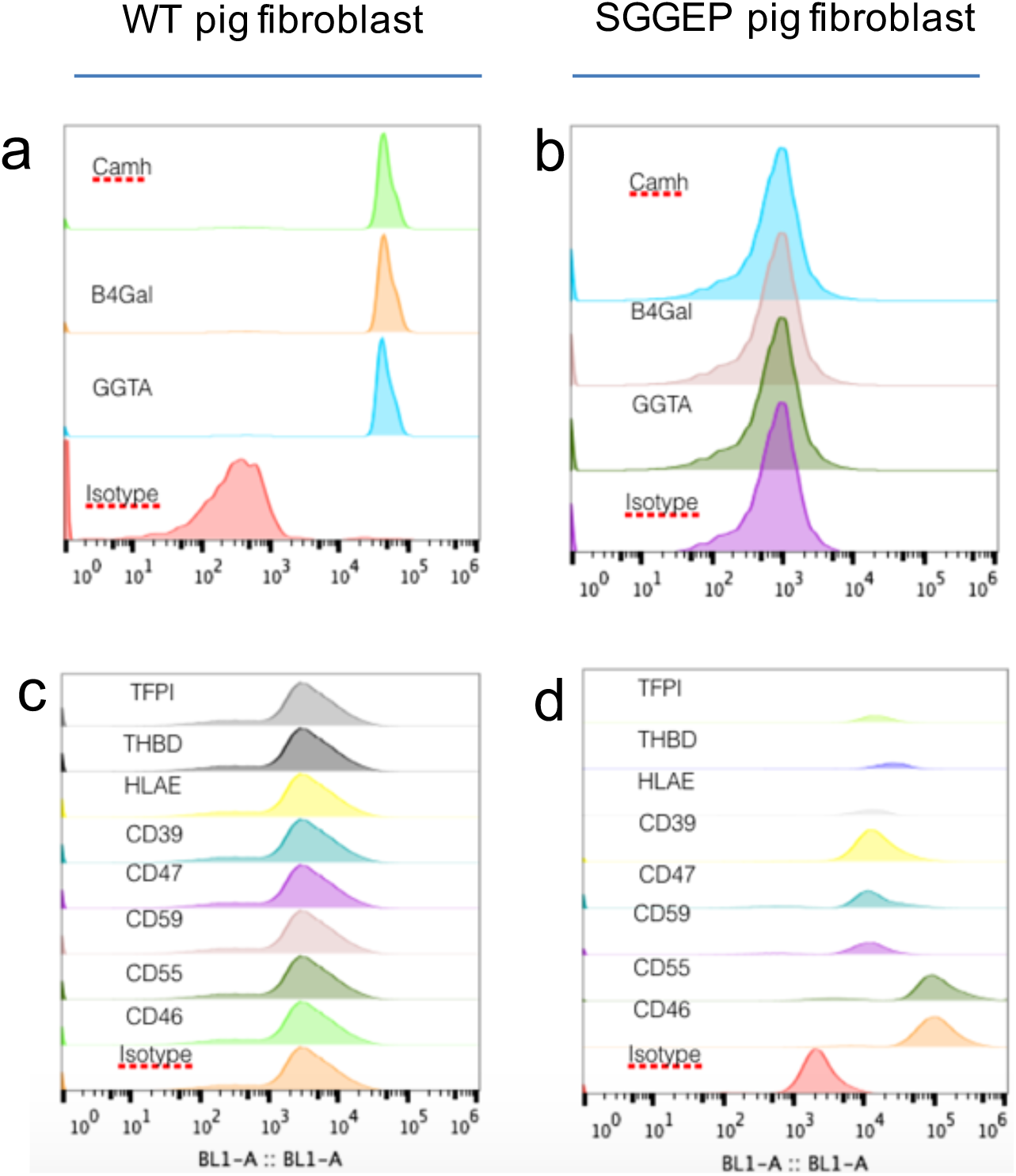
Flow cytometric analysis of WT and SGGEP pig skin fibroblasts phenotype. **(a**) WT pig fibroblasts were stained with the xenoantigens related antibody of GGTA, B4Gal and Camh, xenoantigens binding WT fibroblasts was noted by the marked increase in fluorescence compared to non-specific Isotype antibody stained cells.(b)SGGEP pig fibroblasts were stained with the xenoantigens related antibody of GGTA, B4Gal and Camh, xenogens binding WT fibroblasts was noted by the no increase in fluorescence compared to Isotype antibody stained cells.(c) WT pig fibroblasts were stained with the human proteins related antibodies: from top TFPI,THBD,HLAE,CD39,CD47,CD59,CD55,CD46, no increase in fluorescence in the cells compared to isotype antibody stained cells indicating no expression of these protein in WT fibroblasts. (d) SGGEP pig fibroblasts were stained with the human proteins related antibodies: from top TFPI,THBD,HLAE,CD39,CD47,CD59,CD55,CD46, increase in fluorescence staining in the cells compared to Isotype antibody stained cells indicating expression of these proteins.

Next, we assessed the expression of the eight human proteins in SGGEP fibroblasts (TFPI, THBD, HLAE, CD39, CD47, CD59, CD55, and CD46). Human protein expression in the SGGEP fibroblast was compared to WT pig fibroblast by flow cytometry. WT pig fibroblasts did not express the eight human proteins (Fig. 1c). The increased fluorescence intensity of the eight human proteins in SGGEP fibroblasts indicates successful knock-in of the human genes (Fig. 1d). Taken together, the SGGEP was successfully established, with three xenoantigens deleted and eight human genes stably expressed (see supplemental Fig.S2, supplemental Video1).

### SGGEP skin fibroblasts exhibited the human immunity evasion in vitro

After successfully generating the multiple genes modified SGGEP, we assessed the immunological compatibility of the SGGEP pig to human in vitro. First, SGGEP and WT pig fibroblasts were incubated with pooled human serum (containing preformed IgM and IgG) to determine cross-reactivity with human immunoglobulins. Serum samples from 60 healthy peoples were pooled together for this experiment. Compared to WT pig fibroblasts, SGGEP fibroblasts showed remarkably low binding to human IgM and IgG, suggesting a low possibility for hyperacute immune-rejection in the future xenotransplantation for a variety of Organs (Fig.2a).

Subsequently, we performed a human complement toxicity assay to analyze the immunological response for the SGGEP’s genetic modification. SGGEP and WT pig fibroblasts were incubated with human sera complement (50%) and stained with Propidium Iodide (PI) dye, a cell death marker. Human fibroblasts served as the control. Results showed decreased PI fluorescence in SGGEP fibroblasts compared to WT fibroblasts, indicating increased protection to human complement killing for SGGEP fibroblasts (Fig. 2b)

**Figue2:**
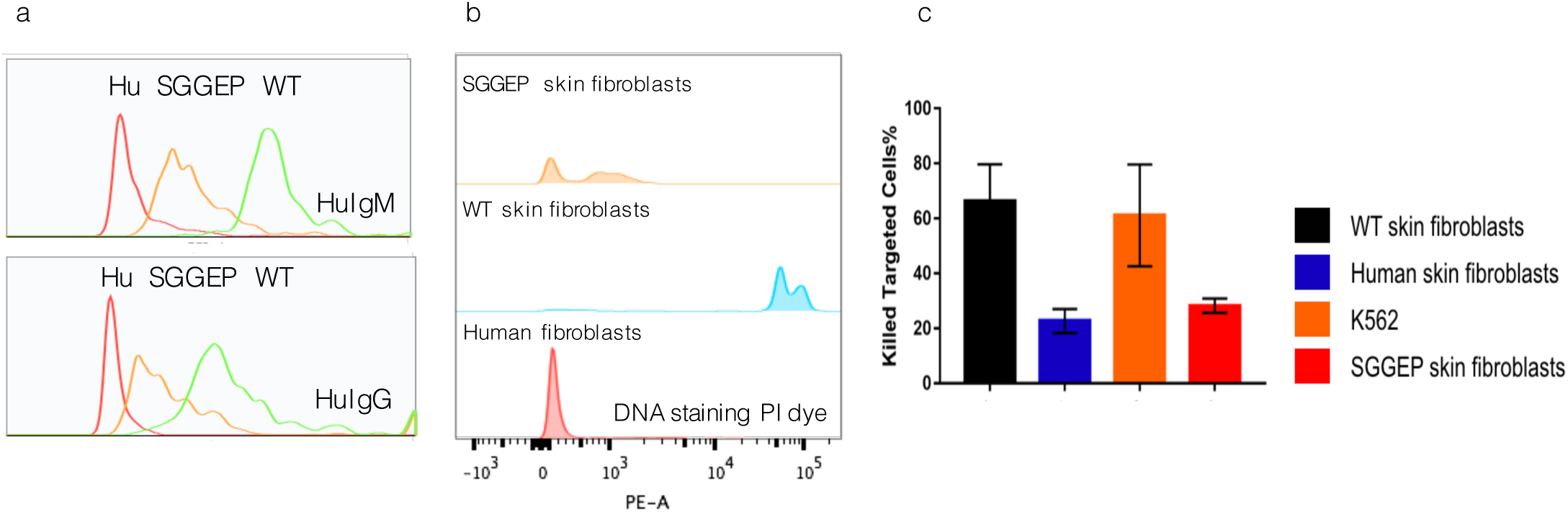
Attenuated human preformed IgG/IgM antibodies binding; human complements killing and NK cells cytotoxicity for SGGEP fibroblasts in vitro. (a)Pooled human serum antibodies binding assays:Skin fibroblasts from WT,SGGEP and human healthy volunteer were incubated with pooled 60 healthy sera (50%), second fluorescent antibodies were applied to show the degree of IgM or IgG binding. Top panel indicated the IgM binding with WT skin fibroblasts is highest, while its binding with SGGEP is much low comparing to that of WT ones and very close to that of human fibroblasts. The low panel showed the same pattern for human IgG binding. (b)Human complement toxicity assays:Skin fibroblasts from WT,SGGEP and human healthy volunteer were incubated withhuman sera with complement(40%) and DNA staining dye PI was applied at the same time. The WT skin fibroblasts showed highest PI fluorescence, indicated highest complement toxicity; human skin fibroblasts did not activate complement showed lowest fluorescence. The SGGEP skin fibroblasts’s fluorescence is close to human one indicated the resistance to human complement killing(c)Natural Killer assay:Skin fibroblasts from WT,SGGEP, human healthy volunteer, and human tumor cell(K562,positive control) were incubated with human Natural Killer cells while DNA staining dye PI was applied at the same time. The WT skin fibroblasts and K562 activated the natural killer cells and the cell membrane was damaged so they exhibited high level of fluorescence. But the SGGEP skin cells showed natural killer cells toxicity resistance. They had much lower level of fluorescence and were very close to that of healthy volunteer.

Finally, we performed a human Natural Killer (NK) cell toxicity assay. SGGEP and WT pig fibroblasts were incubated with human NK cells as well as K562 cells (human cancer cell line) and stained with PI. Human fibroblasts served as the control. WT pig fibroblasts and K562 had an increased proportion of dead cells, whereas the SGGEP fibroblasts had a low proportion of dead cells, comparable to human fibroblasts, indicating protection to NK cell cytotoxicity (Fig. 2c).

As a whole, these results suggest that the multiple genes modified SGGEP we generated has enhanced compatibility with the human immune system and attenuates human antibody binding, complement toxicity, and NK cell cytotoxicity.

### Xenogeneic skin grafts from SGGEP exhibit comparable or longer survival to allografts

After successfully creation the SGGEP with enhanced human immunological compatibility, we next conducted an *in vivo* skin graft study to assess functionality in a non-human primate (NHP) model. Four cynomolgus monkeys received a combination of autologous, allogeneic, SGGEP xenogeneic split-thickness skin grafts, and ADM (Acellular Dermal Matrix) on separate full-thickness dorsal wounds (Fig. 3). All wounds and skin grafts were standardized to 3×3 cm. Autologous skin graft could offer complete clinical healing and permanent closure of wounds (Fig. 3, top panel). Without immunosuppressants, allogeneic skin graft could provide vascularizing comparable to autologous grafts and offer a healed wound environment (Fig.3 the second lane). Rejection occurred from Day14, allogeneic skin graft could survive by day 21 and complete rejection by day 25. SGGEP skin graft could achieve re-vascularization and could offer comparable healed wound environment, barrier and protective function as the allogeneic graft. It underwent rejection from day 14 and could survive by day 25 (Fig.3 the third lane). This clinical appearance was confirmed by histology (Fig.4). Early engraftment and revascularization were observed (Fig. 4b, 4c), progression to rejection by day 25 after transplantation was evidenced (Fig.4d, 4e) with inflammatory cells infiltration and graft loss. By day 29 the grafts were almost completely rejected (Fig.3,the third lane,Fig.4f) by histology. ADM could cover a wound, but it could not achieve a healing environment and could only provide a physical barrier within 7days. By 7 days it contracted and began to peel off by day 14 (Fig.3 the fourth lane).

**Figue4:**
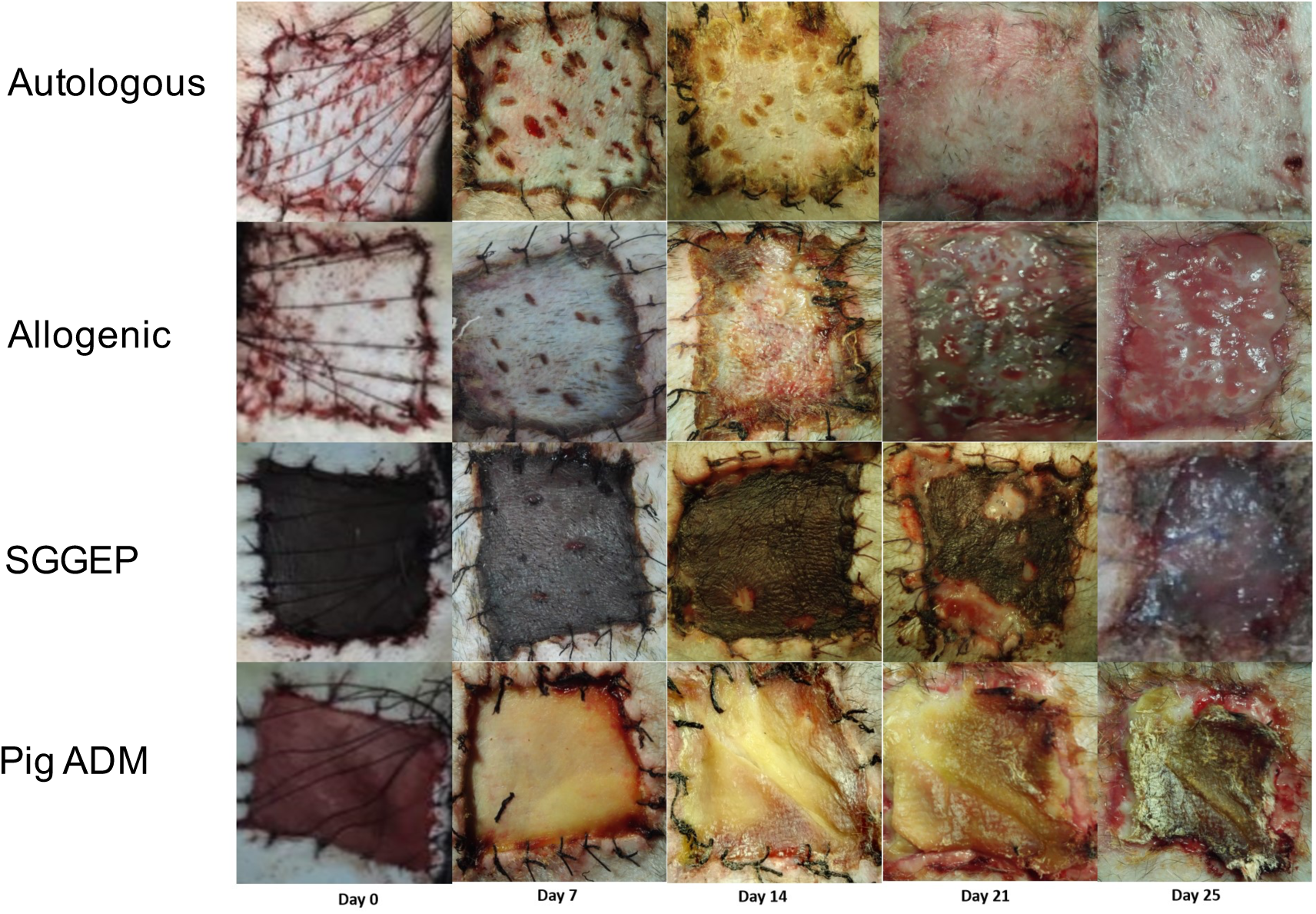
Time course of split-thickness skin graft gross pathology appearance. Autologous skin graft could offer complete and permanent closure of wounds. Without immunosuppressants Allogeneic skin graft could provide revascularizing comparable to autologous grafts and offer a healed wound environment. It occurred rejection form Day 14, could survive on day 21 and complete rejection on day 25. SGGEP skin graft could achieve re-vascularization and could offer comparable healed wound environment, barrier and protective function as allogeneic graft. It experienced rejection from day 14 and could survive on day 25.Pig ADM(acellular dermal matrix) could close wound, but could not achieve a healing environment and could only provide physical barrier within 7days.After 7days it contracted and was began to peel off by day 14.

**Figure 4.**
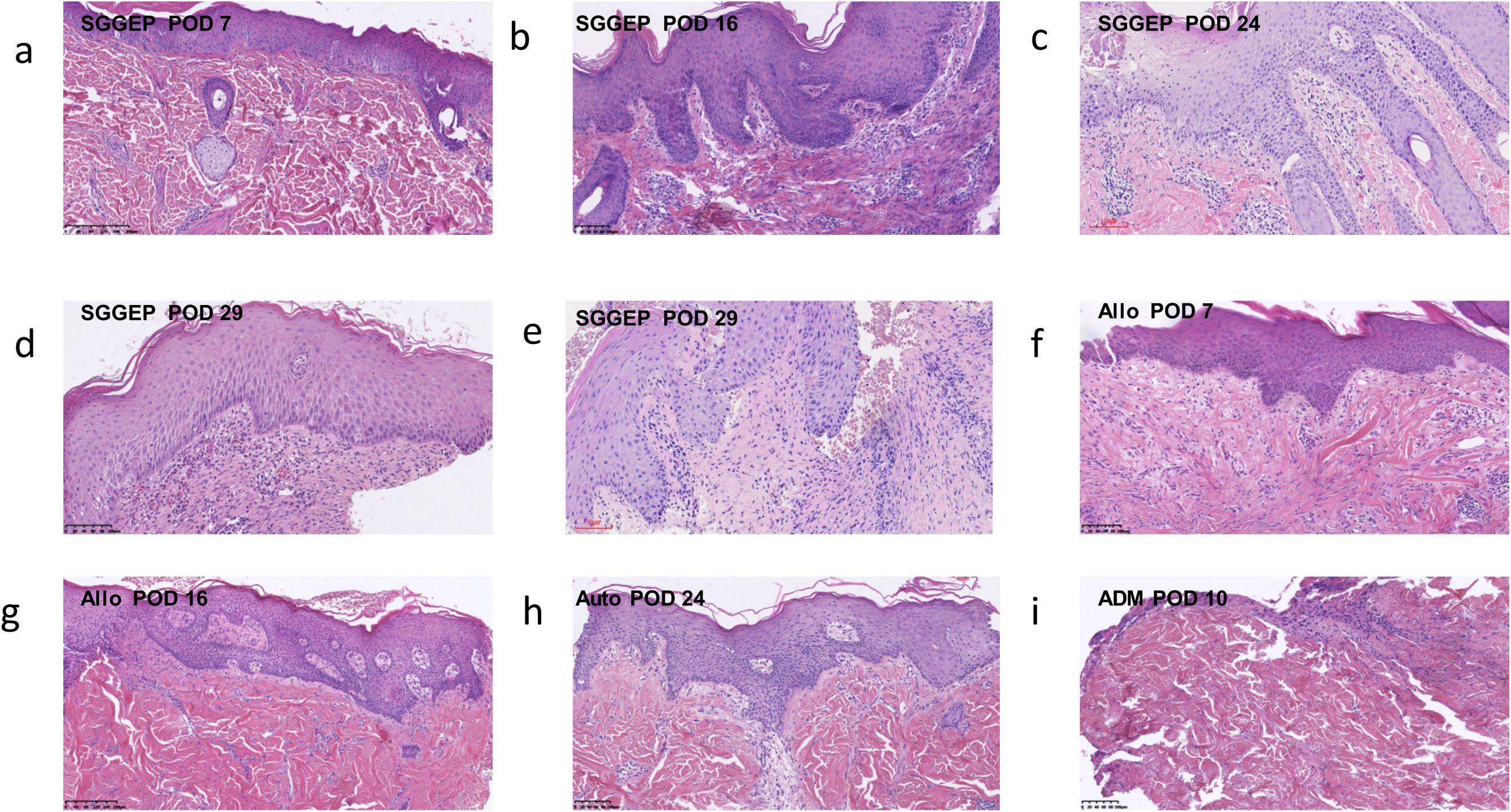
The representative images of histology HE staining were showed from biopsies after skin xenotransplantation at indicated time points. (a)SGGEP graft on Post-Operation Day 1(POD1),(b) SGGEP graft on POD 16,(c) SGGEP graft on POD24,(d) SGGEP graft #1on POD29, (e) SGGEP graft #2 on POD29,(f) Allogenic graft on POD7,(g) Allogeneic graft on POD 16,(h) Autologous graft on POD24,(i) ADM

In summary, our *in vivo* xeno-skin graft study showed enhanced immunological compatibility with NHP, consistent with our *in vitro* studies. These findings provide greater evidence and clinical confidence to support a human clinical trial with SGGEP organs.

## Materials & Methods

### Generation and selection of SGGEP(3KO+8KI) pig skin fibroblasts

The SGGEP pig was created as demonstrated in Figure S1The triple KO pig fibroblasts were generated according to methods discussed in the current literature (9, 10). Specific gRNA sequence for GGTA is GTC GTGACCATAACCAGA; for CMAH is AGAAACTCCTGAA CTACA; for B4GAL is AGGAAAGCTATAACTTGG.

Primary skin-derived fibroblasts were simultaneously transfected with three Cas9 endonucleases (Thermofisher, USA), gRNA complex (true guide synthetic gRNA, Thermofisher, USA) specific for the GGTA, CMAH, and B4Gal genes with the Neon Transfection System (Life Technologies, USA). GGTA negative cells were selected by counter-selection using streptavidin-coated magnetic beads (Dynabeads, Life Technologies, USA) and biotin-conjugated isolectin B4 (Enzo Life Science, USA) using a magnetic field. Cells without GGTA flowed out of the column due to a lack of carbohydrate xenoantigen. Triple knockout cells were isolated by fluorescence-activated cell sorting. These cells do not bind the DBA lectin, which targets carbohydrate structures produced by the B4Gal enzyme, as well as poly-clonal chicken anti-Neu5GC antibody (Biolegend, USA) plus secondary staining with FITC-labelled donkey anti-chicken IgY (Jackson ImmunoResearch, USA). Phenotypic selection of cells without GGTA, B4Gal and CMAH expression were used for further genome edition. The human proteins were assigned to 3 Piggybac vectors (System Biosciences, USA) by Golden Gate Assembly method (11, 12). Vector 1 has the CD46, 55 and 59; vector 2 contained the CD39, THBD, TFPI; the vector 3 bears CD47and human HLA-E. All the vectors were driven by promoter Human elongation factor-1 alpha (EF-1α). The triple knockout fibroblasts then were cultured for two weeks, collected, and then transfected with three piggyBac vectors expressing multiple human proteins with the Neon Transfection System with the presence of piggyBac Transposase (System Biosciences, USA). Two days after, successfully transfected CD55, CD47 and CD39 positive cells were selected by Fluorescence-activated Cell sorting (FACS) and then used for somatic cell nuclear transfer to generate SGGEP.

### Nuclear Transfer and Embryo Transfer

Pig ovaries were abstained from a local slaughterhouse and shipped to the laboratory within one hour in 0.9% saline maintained at 37 °C. The maturation of oocytes, enucleation, microinjection, and the fusion of reconstructed oocytes in vitro were conducted following previously described methods (13, 14). The reconstructed embryos were cultured in porcine zygote medium in 5% CO2 at 39 °C for 14–16 hours, then the embryos in good shape were surgically transferred into the oviduct of a surrogate. Naturally, the surrogates gave birth to the cloned piglet, named as Rescuer (See Supplement Fig.S2, Supplemental video1).

### Flow cytometric phenotyping of SGGEP skin fibroblasts

Xenoantigen expression was detected by methods discussed in the literature (5, 9). The Neu5Gc kit was obtained from BioLegend, USA. B4Gal phenotype was analyzed with Dolichos biflorus agglutinin (DBA lectin), stained with fluorescein (Vector Laboratories, USA). All cells were suspended in HBSS (2×10^6^cells/mL) with Neu5Gc free blocking buffer for 15 minutes and then incubated for 30 minutes at room temperature with the appropriate lectin or antibody. Cells were then washed with blocking buffer. All flow cytometry analyses were performed on the Sony SA3800 Spectral analyzer. The following provides a brief description of the protocol: 1) Harvest and wash the cells (single-cell suspension), 2) adjust the concentration to 1-5×10^6^ cells/mL in ice-cold FACS buffer (5-10% FBS) using polystyrene round-bottom 12 × 75 mm BD Falcon tubes, 3) add 0.1-10 μg/ml of the primary labelled antibodies to100 μl of cell suspension in each tube, 4) incubate cells for 30 min at room temperature in the dark, 5) wash the cells three times by centrifugation at 1500 rpm for 5 minutes and resuspend in 200 μl FACS Buffer, and 6) Analyze the cells by flow cytometry. Results were managed with Flowjo V10 software. Anti-Human CD55, CD59, CD46, CD47, CD39, and FITC conjugated antibodies were purchased from BD, USA. The THBD-488 conjugated antibody was from Abcam, USA. The HLAE and TFPI’s primary antibodies were from LSBio, and the second FITC antibodies were from BD, USA.

### Human antibody binding to SGGEP skin fibroblasts

Sera from over 60 healthy volunteers were purchased from Innovative Research, USA, and heat-inactivated at 57 °C for 30 min. Cells stained with goat anti-human IgG Alexa Fluor 647 (Life Technologies, USA) and Donkey anti-human IgM Alexa FITC (Southern Biotech, USA) were selected.The cells of 5×10^5^ were incubated with a 50% mixture of serum at 4°C for 30 minutes, then washed three times and stained with anti-IgG or IgM antibodies as described. Cells were washed again and analyzed by flow cytometry.

### Human complements toxicity assay for SGGEP skin fibroblasts

The cells of 5×10^5^ were incubated with 50% human complements and stained with PI at 37°C for 45minutes. The cells were then washed again and analyzed by flow cytometry.

### Human Natural Killer cells toxicity assay for SGGEP skin fibroblasts

Human natural killer cells (Stemcell Technologies, Canada) were incubated with 5×10^5^ skin fibroblasts for 60 min at 37°C, at an effector: target=10:1, and stained with PI. Cells were washed again and analyzed by flow cytometry. Anti-CD56-647 conjugated antibody (BD, USA) served as a marker of natural killer cells.

### Animals

All studies were approved by the First Affiliated Hospital of Nanchang University Institutional Animal Care and Use Committee (IACUC) and performed in accordance with the Guidelines for the Humane Treatment of Laboratory Animals (2006) issued by the Chinese Government.

(http://www.most.gov.cn/fggw/zfwj/zfwj2006/200609/t20060930_54389.htm. Cynomolgus monkeys, between 4-6 years and weighing 4.2-4.5kg, were obtained from the Guangdong Landau Biotechnology Co. Ltd. All animals were undergone through routine pathogen screening and quarantine prior to the start of studies.

### SGGEP Skin graft harvest and skin graft transplant

Four cynomolgus monkey recipients were anesthetized with 2.5mg/kg diazepam IM and 10mg/kg ketamine IM. Then, endotracheal intubation was performed under anesthesia (2% isoflurane and oxygen). The dorsum was shaved using clippers. 3 × 3cm full-thickness wounds were then made by excision of the skin. The incision was made by the scalpel until the dorsal muscle to remove the subcutaneous tissue and fascia. Saline soaked gauze was applied to stop bleeding.

Swine donors were treated, intubated the same way with anesthesia. Cynomolgus monkeys were kept under general anesthesia in the prone position. The pigs’ skin was disinfected with Povidone-Iodine and rubbed with 70% ethanol. Split-thickness skin grafts were harvested from Cynomolgus monkey or pig with a dermatome(Humeca, Netherland)set depth of 0.4 mm. Skin grafts for immediate use were stored at 4°C.

Skin grafts from pigs were sutured to full-thickness wound bed in Cynomolgus with 3–0 Ethilon by interrupted sutures. Pressure bandaging was achieved and attached firmly by protective jackets. Ceftriaxone sodium(80mg/kg)IV was administered once before and once after the operation. The rejection was defined as a 90% loss of the original graft in this study. Pictures were taken to document the gross appearance of skin grafts every 4 days. A 3mm needle biopsy was taken at the indicated time for histological assessment.

## Discussion

Organ transplantation is the only curative solution for organ failure. Yet, the worldwide organ shortage limits the number of life-saving procedures possible. xenotransplantation is an alternative solution to this problem. The pig model is widely used in biomedical research. Compared to NHP and other species, the miniature pig is the most ideal species for xenotransplantation. However, pigs are not phylogenetically close to humans. Thus, immune rejection and molecular incompatibility can occur in xenotransplantation. Thanks to the advancement in genetic engineering, especially CRISPR/Cas9, the modification of genetic material in pigs has become possible. As a result, considerable progress has been made in xenotransplantation of major organs using genetically modified pigs.

It is well known that the three-known carbohydrate xenoantigates (GGTA, B4gal, and CAMH) can trigger hyperacute immune rejection. Various NHP studies have shown that the deletion of these xenoantigens in pig organs can extend graft survival in NHP (7). Studies have also reported that the additional expression of certain human transgenic proteins may be beneficial for xenografts from pig to human, such as human complement-regulatory proteins (hCRPs) (CD46, CD55, CD59), human coagulation-regulatory proteins thrombomodulin (THBD), tissue factor pathway inhibitor (TFPI), CD39, CD47and HLA-E (7, 15). The generation of multi-gene modified pigs with xenoantigen deletions and human protein expression is a major scientific advancement.With such kind of the pig, it is likely that heart, kidney and pancreatic islets xenotransplantation clinical trials would be launched in the near future (7, 15).

Our objective in this study was to generate a genetically modified donor pig that can serve as a potential organ source for future clinical transplantation. We utilized CRISPR/Cas9 to create triple xenoantigen KO pig cells. Three enzymes involved in manufacturing xenoantigens were knocked out to generate a GGTA/ B4GAL/CMAH KO pig. We then expressed critical human proteins including human complement system negative regulatory proteins (CD46, CD55, and CD59); human coagulation system negative regulatory proteins thrombomodulin (THBD); tissue factor pathway inhibitor (TFPI); CD39; macrophage negative regulatory proteins (human CD47); and natural killer cell negative regulatory human HLA-E in the GGTA/ B4GAL/CMAH KO cells to generate a new smart specific gene-modified pig by nuclear transfer. In vitro studies by flow cytometric analysis of the wild type and SGGEP skin fibroblasts confirmed our SGGEP lacks three xenoantigen expression with our expected additional molecule protein expression. Analysis of the SGGEP’s genetic modifications on the immunological response showed attenuated human antibody binding, complement toxicity, and NK cell cytotoxicity compared to WT pig cells, suggesting enhanced compatibility with the human immune system.

We then further assessed the functional impact of our SGGEP *in vivo* in NHP. Four cynomolgus monkeys received a combination of autologous, allogeneic, SGGEP xenogeneic split-thickness skin grafts and ADM on separate full-thickness dorsal wounds. In this study, the SGGEP skin graft could remain engrafted up to 25 days. The *in vivo* studies showed that xenogeneic skin grafts from SGGEP exhibit comparable or longer survival to allografts. Therefore, our *in vivo* skin graft study demonstrated enhanced immunological compatibility with NHP, consistent with the *in vitro* findings.

Burn injuries can be devastating and severely debilitating, making such injuries a major global health concern (16, 17). Autografts are considered the optimal way in cases of deep second- and third-degree burn wounds; however, tissue source is limited in availability, especially for large burns. In this scenario, allogeneic cadaver skin grafts are an appropriate alternative; however, the high cost, low availability and possibility of transmitting pathogens limit the clinical use of this procedure. In 2018, the U.S. FDA approved the first human clinical trial for the topical application of live skin from GGTA single KO pig for the treatment of severe burns, which is a critical milestone for xenotransplantation progress. Our *in vivo* study showed that SGGEP skin grafts survived functionally longer on NHP than GGTA single KO pig skin (18). SGGEP skin grafts are functionally comparable to allogeneic skin grafts. These grafts can provide barrier protection and a healing environment for full-thickness wounds to heal. Thus, the SGGEP skin grafts are an appropriate therapeutic alternative for cadaver allogeneic skin grafts in patients with large burns, especially in trauma and emergencies.

However, it is important to note that extensive genome editing in pig somatic cells is not practical due to the limitation of telomere length. Currently, the ideal cell for genetic modifications such as porcine ES or iPS is not available. Although porcine ES or iPS like cells were reported, these ES or iPS like cells cannot give rise to a live pig so far and it is very difficult for them to become donor cells for nuclear transfer (19-21). With limited options, primary pig somatic cells were used as the starting material for the vast majority of pig genome genetic engineering. Complex and extensive editing requires long cell culture periods and frequent passaging that could unavoidably exceed telomere length and cause cell ageing, apoptosis, cell cycle arrest or death.

It should be also mentioned that another potential risk in xenotransplantation is a porcine endogenous retrovirus (PERV) which is integrated into pigs’ genome. However, the risk of the pathogenic effect is considered very low (22-24). In addition, numerous PERV infection experiments, as well as preclinical and clinical xenotransplantation trials such as islet xenotransplantation trials, have failed to show transmission (25, 26). There are also several antiviral medications on the market that may be used to treat potential transfer-related clinical risk. Moreover, deleting all copy numbers may be harmful to the pig cell, restricting the number of other necessary genetic modifications that can be made (7). Thus, the low risk of PERV transmission and pathogenicity should not hinder launching a clinical trial, US FDA approval of GGTA knockout pig for skin xenotransplantation clinical trial in 2018 speaks for this. Therefore, upon carefully weighting harm and benefit, we employed a strategy to create such SGGEP which is consistent with the current scientific community. Our combination of the multi-gene modified pig will certainly provide a tool to accelerate xenotransplantation progress.

In summary, we generated an SGGEP with a better immunological and molecular compatibility profile for humans. And we determined a good combination of gene modifications for clinical application. Our *in vitro* and NHP *in vivo* study provides great evidence and clinical confidence to support a human clinical trial with SGGEP organs. With promising research from the scientific community and generation of our SGGEP pig, the next step is to launch investigator-initiated trials to confirm safety and efficacy in humans. This technology will offer an attractive alternative in the management of patients with severe burns. The skin grafts can be cryopreserved, stored for unexpected catastrophic events, and transported worldwide for global use (Fig. 5). Eventually, we may be able to provide patients with severe burns with lifesaving treatment. As the skin is considered the vital, unique and immunogenicity organ, our preliminary success in skin xenotransplantation using the combination of multi-gene modified pig in NHP provides the approval of the concept, paves a way to initiate the other organ preclinical trial and clinical trial, implies a success of these organs’ xenotransplantation.

**Figure 5.**
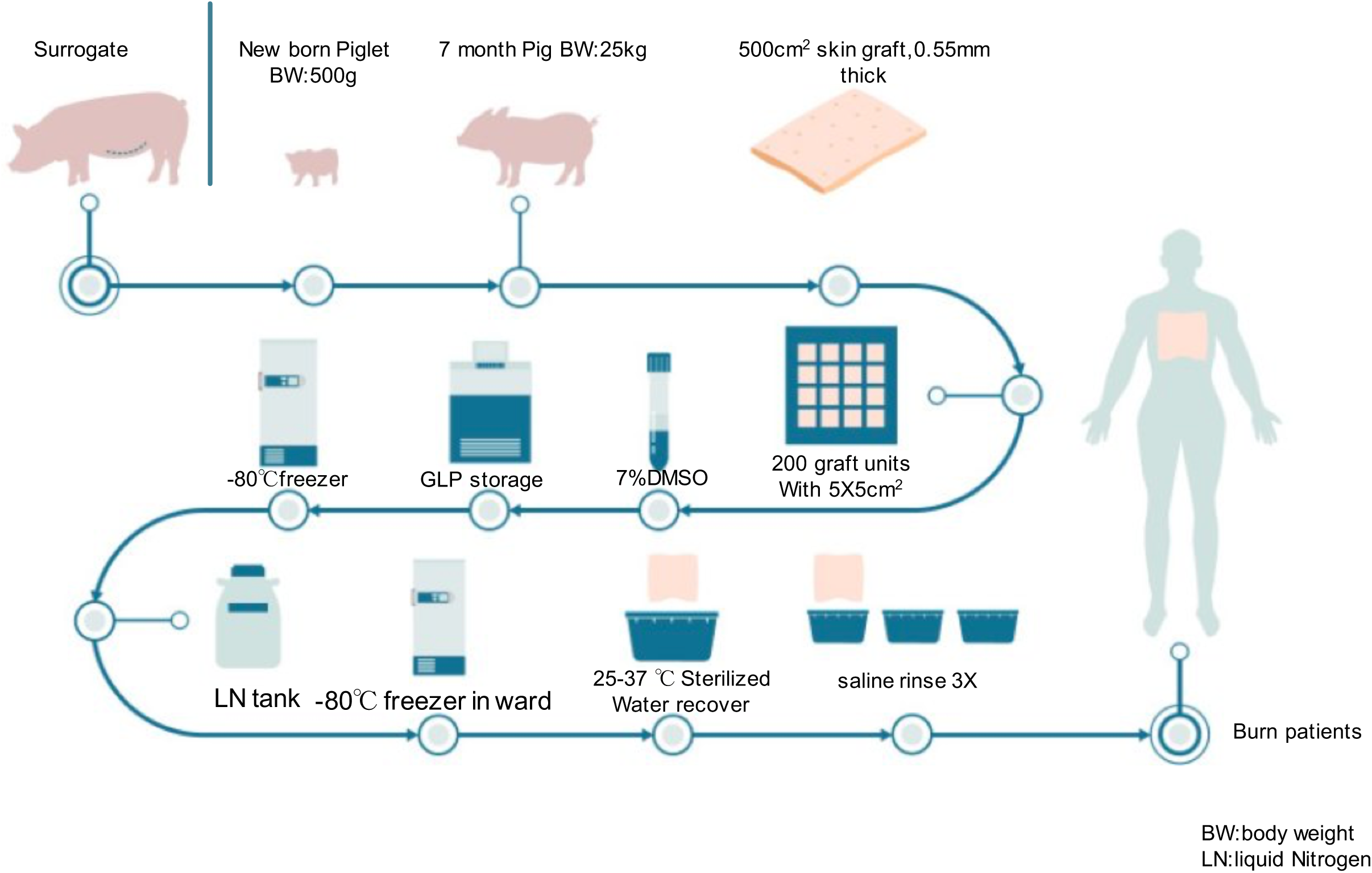
SGGEP skin xenotransplantation working model.

## Acknowledgements

L.Z., Y.Z. was supported by the grant Natural Science Foundation of China(NSFC 81660364 and NSFC 81760343). X.Z. was funded by grants Natural Science Foundation of Guangdong Province (2017A030308004) and Natural Science Foundation of Guangzhou City (201804020011). G.Z. was supported by the National Key R&D Program of China (2017FA105202) and Shenzhen Scientific & Innovation Committee (GJHZ20180928155604671)

