## Supplemental Figures and video for "Selective germline genome edited pigs and their long immune tolerance in Non Human Primates"

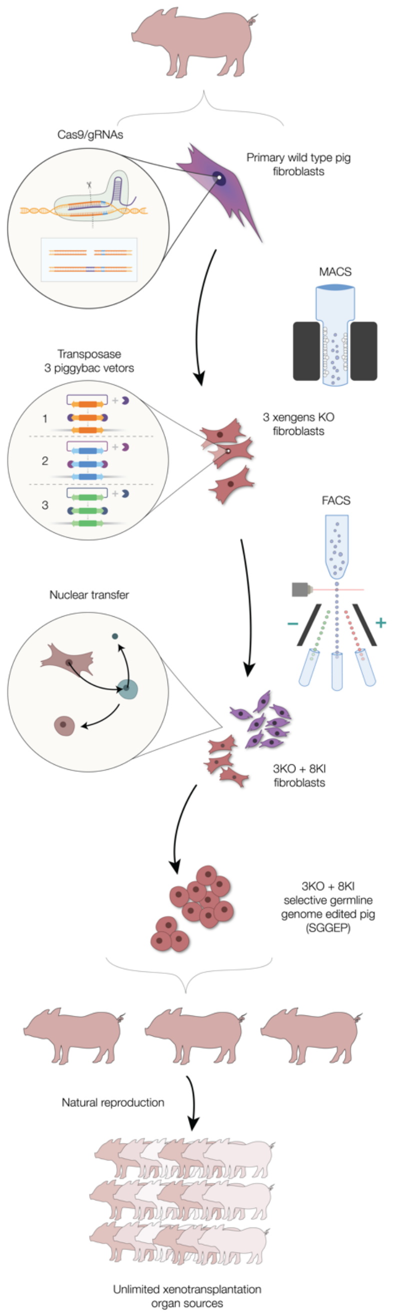


MACS:Magnetic-activated cell sorting

FACS:Fluorescence activated cell sorting

**FigureS1 The flowchart for generation of SGGEP**

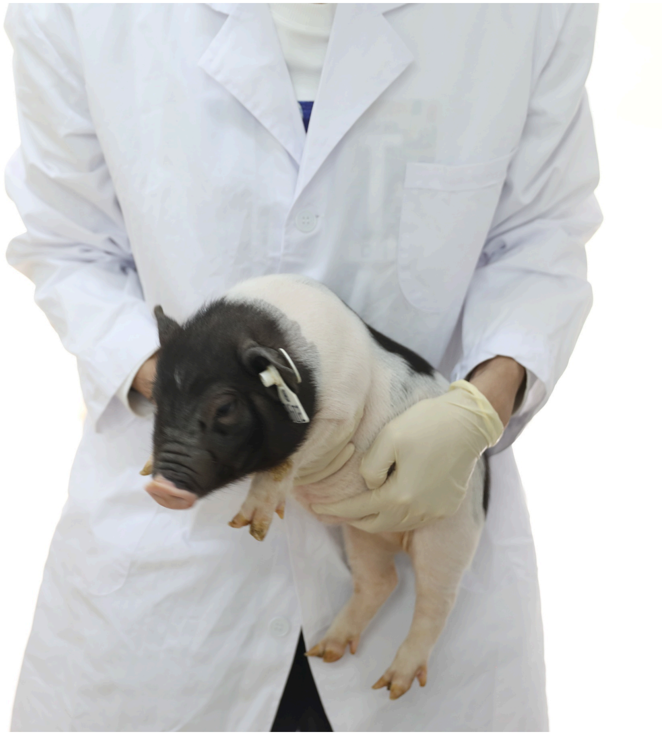

FigureS2 SGGEP- Rescuer

### Supplemental video1

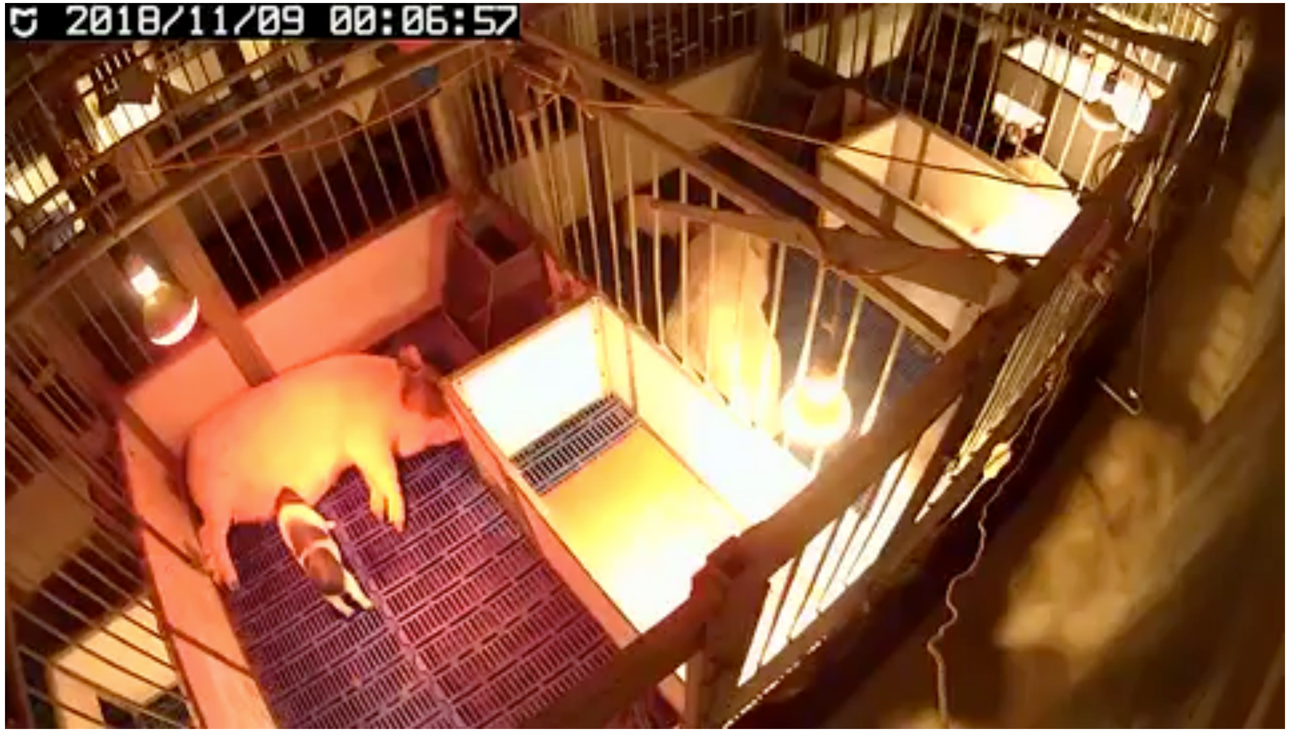


**SGGEP-Rescuer**
